# A High-Quality Genome Assembly of *Chaetoceros muelleri* Reveals Extensive Gene Duplication, Functional Diversification, and Unique Lineage-Specific Innovation

**DOI:** 10.64898/2026.04.04.716509

**Authors:** Anushree Sanyal, Elinor Andrén, Christian Tellgren-Roth

## Abstract

Diatoms are major contributors to marine primary production, yet high-quality nuclear genome resources remain scarce for ecologically dominant lineages such as *Chaetoceros*. Here, we present the first high-quality nuclear genome assembly of *Chaetoceros muelleri*, generated from living cells resurrected from resting spores preserved in Baltic Sea sediments and sequenced using PacBio HiFi long-read technology. The assembly is compact (43□Mb), highly contiguous (N50□=□1.40□Mb), and highly complete (93% BUSCO).

Comparative analyses across 14 diatom genomes revealed extensive lineage-specific and expanded gene families in *C. muelleri*, alongside a small, conserved core genome, reflecting rapid evolutionary turnover. Functional enrichment highlighted diversification of polysaccharide biosynthesis, vesicle-mediated trafficking, membrane remodelling, and transcriptional regulation, consistent with adaptations linked to frustule formation and environmental responsiveness. Transposable elements (TEs) strongly shape the genome, accounting for ∼18% of the assembly, with dominant LTR retrotransposons and a large fraction of unclassified repeats suggesting novel or highly diverged TE lineages. Enrichment of DNA replication, recombination, and repair functions further indicates compensatory genome maintenance associated with TE-driven structural dynamics.

Direct comparison with *C. tenuissimus* revealed contrasting patterns of gene family expansion and regulatory innovation, underscoring divergent evolutionary strategies within *Chaetoceros*. By integrating resurrection ecology with long-read genomics, this study provides a foundational genomic resource for *C. muelleri* and highlights the role of TE-mediated genome plasticity in diatom evolution.

## Introduction

Diatoms (Bacillariophyta) are among the most diverse and ecologically important groups of photosynthetic protists, contributing an estimated 40 % to marine primary production and 20–30% of global carbon fixation and playing central roles in marine biogeochemical cycles, global carbon export (Field et al. 1998, Mann & Droop 1996; Malviya et al. 2016, Tréguer et al. 2018), and are bioindicators (Benoiston et al. 2017). The diatom genus *Chaetoceros* is the largest and most speciose with more than 500 described species spanning polar, temperate, and tropical waters and dominating plankton communities in many coastal and upwelling systems (Nelson et al. 1995; Guiry & Guiry 2020). *Chaetoceros* species significantly influence global carbon cycling, ecosystem functioning, and marine food webs (Zigic et al. 2026).

*Chaetoceros* species exhibit extensive morphological diversity, rapid speciation, high environmental responsiveness, and strong population structure in regions undergoing climate-driven change (Nef et al. 2022, Vrana et□al.□2023). *Chaetoceros muelleri* represents a particularly valuable species for genomic investigation due to its ability to form long-lived resting stages and its ecological relevance in marine systems. It is ubiquitous, ecologically versatile, occurring in both marine and brackish systems (De□Luca et□al.□2019), dominates phytoplankton communities during nutrient-replete conditions (Lovio-Fragoso et□al.□2019), and is widely used in aquaculture due to its high lipid productivity and nutritional quality (Batista et al. 2015, Naghdi et al. 2016).

Despite its ecological and economic importance, no high-quality nuclear reference genome has been available until now, limiting the study of its evolutionary biology, ecological adaptation, and genomic innovations. The genus is represented largely by organellar genomes. Chloroplast genomes of seven *Chaetoceros* species (Xu et al. 2021), including *C. muelleri* were recently sequenced (Li & Deng 2021). Also metagenome-assembled genomes from Tara Oceans surveys revealed strong environmental selection and nutrient-driven genomic divergence in natural *Chaetoceros* populations (Nef et al. 2022). The characterization of its organellar genome underscored its phylogenetic positioning within the genus (Li & Deng 2021), but provided little insight into nuclear gene content, transposable element (TE) dynamics, or metabolic pathway diversification. Nuclear genomes are essential for identifying the genetic bases of these processes, resolving gene family evolution, quantifying TE impacts, and understanding the molecular mechanisms shaping frustule formation, nutrient acquisition, and environmental resilience. However, high-quality, long-read nuclear genome assemblies for *Chaetoceros* species are still extremely limited, and genomes of only two species *Chaetoceros tenuissimus* and *Chaetoceros gracilis* (Hongo et al. 2021, Kumazawa et al. 2022) have been sequenced.

Sequencing the *C. muelleri* nuclear genome therefore fills a critical gap in diatom genomics. It expands the limited set of centric diatom genomes—currently dominated by *Thalassiosira* and *Skeletonema* species (Ambrust et al. 2004, Lommer et al. 2012, Liu & Chen 2024a, 2024b, Di Costanzo et al. 2025, Nagai et al. 2025),and adds a key representative of the ecologically dominant *Chaetoceros* lineage. A high-contiguity, long-read assembly enables deeper interrogation of gene family evolution, repeat-driven genome diversification, and lineage-specific innovations underlying ecological success. Furthermore, including *C. muelleri* within comparative frameworks spanning centric and pennate diatoms will enhance our understanding of the evolutionary trajectories shaping diatom diversity, functional adaptation, and genomic plasticity.

Diatom resting spores recovered from marine sediments provide an extraordinary opportunity to directly access historical phytoplankton genotypes and reconstruct evolutionary trajectories across ecologically meaningful timescales. Resting stages of diatoms are known to remain viable for exceptionally long periods, with demonstrated survival from centuries to several millennia under anoxic sediment conditions, retaining enough cellular integrity to resume growth when re-exposed to light and oxygen (Härnström et al. 2011, Sanyal et al. 2022, Bolius et al. 2025). Unlike sedimentary ancient DNA—which is typically fragmented and chemically degraded—resurrected resting spores allow researchers to obtain living cells and generate high-quality genomes that accurately represent historical populations. Resting spores act as natural genomic archives, allowing recovery of historical genotypes unaffected by recent culturing or laboratory selection.

Recent studies from the Baltic Sea have revived diatom spores as old as ∼7,000 years (Sanyal et al. 2022, Bolius et al. 2025), showing that dormant stages can be metabolically reactivated and display physiological traits comparable to their contemporary populations (Bolius et al. 2025). The Baltic Sea provides an exceptional setting for this approach as it is a semi-enclosed basin that has experienced extensive nutrient loading, and hypoxia during the past century due to eutrophication and climate-driven physical changes (Meier et al., 2018; Carstensen et al. 2014). The Baltic Sea, with its anoxic sediment layers and high-resolution stratigraphy offers a globally unique setting for such studies due to its accelerated climate change signals and exceptional preservation of resting spores (Carstensen et al. 2014, Conley et al. 2009, Moros et al. 2017, Zillén et al. 2008). Sediment archives from this system thus serve as natural time capsules, preserving genetically intact diatom resting spores from distinct environmental periods. By resurrecting resting stages (resting spores and cells) and sequencing nuclear and organellar genomes from these revived lineages (Härnström et al. 2011, Sanyal et al. 2022, Bolius et al. 2025, Schmidt et al. 2025), direct comparison between historical and contemporary genomes become possible, providing an unparalleled framework for investigating long-term genomic responses to environmental change in one of Earth’s most human-impacted marine ecosystems.

Here, we report the first high-quality nuclear genome assembly of *C. muelleri*, assembled from living cells revived from resting spores preserved in Baltic Sea sediments and sequenced using long-read technology. We characterize its gene content, protein domain architecture, transposable element (TE) landscape, and patterns of species-specific and expanded orthogroups (OGs). We further place *C. muelleri* within a broader phylogenomic context by comparing its genomic features with those of 13 additional centric and pennate diatoms to identify lineage-associated genomic signatures. This work provides a foundational genomic resource for *Chaetoceros* research and offers new insights into diatom evolution, TE-driven genome restructuring, and functional innovation in an ecologically and biotechnologically important lineage.

## MATERIALS AND METHODS

### Sampling

Sediment sampling was carried out from R/V *Electra* at Askö in September 2020 using a 1-m gravity corer to recover the uppermost unconsolidated sediment. A core measuring 85□cm in length was retrieved and stored in a refrigerated container onboard for transport to the laboratory. In the lab, the core was extruded from the bottom and sectioned into 1-cm intervals from the top. Each slice was immediately transferred into labelled zip-lock bags. The sampling station, designated EL20-SH01-01 (57°58.6384′N, 17°57.3703′E), is located at a water depth of 198□m (Fig.□1).

**Fig. 1.**
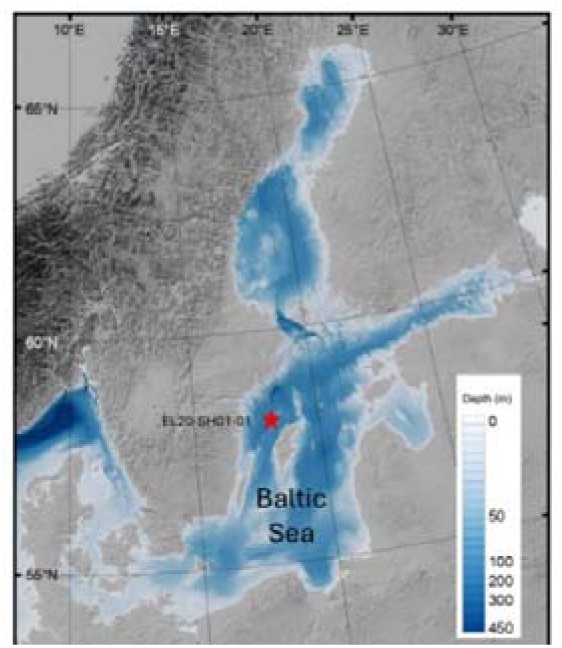
Overview map and bathymetry of the Baltic Sea. The red star indicates the sampling station EL201ZSH011Z01 (57°58.6384′N, 17°57.3703′E), where a short gravity core was collected at a water depth of 198□m.

### Lithology and sample selection

The uppermost section of the core (0–37□cm) consisted of black, gas⍰rich mud with high organic content and no visible laminae. The top 18□cm was very soft, becoming slightly more consolidated below this depth. The lower part of the core, beneath 37□cm, comprised brownish mud with sulphide staining and likewise lacked visible laminae.

Based on the results from radiometric dating and organic matter measurements (loss⍰on⍰ignition), 12 sediment samples from 0 -32 cm corresponding to 2020 to 1965 CE (Table S1) were selected for the revival of *Chaetoceros* resting spores and subsequent cultivation to obtain high⍰quality DNA.

### Radiometric dating and age modelling

Radiometric dating using ^210^Pb and ^137^Cs was performed on sub-samples from the sediment core. Activities of ^210^Pb, ^226^Ra, ^137^Cs and ^241^Am were measured by direct gamma spectrometry at the Environmental Radioactivity Laboratory, University of Liverpool, using Ortec HPGe GWL-series well-type, low-background intrinsic germanium detectors (Appleby et□al.,□1986). ^210^Pb was quantified via its 46.5□keV gamma emission, while ^226^Ra was determined through the 295□keV and 352□keV gamma rays emitted by its daughter isotope ^214^Pb after storing samples for three weeks in sealed containers to allow for radioactive equilibrium. ^137^Cs and ^241^Am activities were measured using their characteristic gamma emissions at 662□keV and 59.5□keV, respectively. Detector efficiencies were calibrated using certified reference sources and sediment standards of known activity. Corrections for self-absorption of low-energy gamma rays were applied following the procedures described in Appleby et□al. (1992). Supported ^210^Pb activity was assumed to be equivalent to the measured ^226^Ra activity, and unsupported ^210^Pb was calculated by subtracting supported 2^210^Pb from total measured ^210^Pb. ^210^Pb dates were calculated using the Constant Rate of Supply (CRS) model (Appleby & Oldfield,□1978). The depths corresponding to the 1986 Chernobyl accident and the 1963 maximum of atmospheric nuclear weapons testing were identified using the ^137^Cs and ^241^Am activity profiles. The final age–depth model was established following an integrated assessment of all available data, applying the procedures described in Appleby (2001) to determine the most robust chronology. The results of the radiometric analyses indicate a relatively uniform sedimentation rate since the early 1990s (Table□S2). Sediments below 33□cm are highly compacted and most likely reflect a hiatus in the sediment record, with material below 39□cm being substantially older.

### Isolation and germination of single resting spores from sediments from the last century

Only the resting spores which germinated were isolated. The single spores were isolated with care following the protocol of Throndsen (1978). Full methodological details regarding the germination and isolation of resurrected resting spores are provided in Sanyal et al. (2022).

### Growing the unialgal Chaetoceros cultures

Unialgal cultures of *C. muelleri* BS20 derived from individually revived resting spores originating from the last century were established following Sanyal et□al. (2022). For an in-depth description of growing the *C. muelleri* cultures, see Sanyal et al. (2022).

### Genome Characterization

#### DNA Isolation from Revived Cultures of Last-Century Diatom Resting Spores

Total DNA from the unialgal *C.*□*muelleri* BS20 culture was extracted using a gentle lysis and phenol–chloroform protocol optimized for diatoms, as described in Sanyal et□al. (2022).

#### PCR and sequencing

PCR amplification and Sanger sequencing were performed to confirm that the taxa is *C. muelleri* using taxonomic markers for centric diatoms used in a previous study by Lee et al. (2013). Comprehensive details regarding PCR amplification and sequencing methodologies are provided in Sanyal et□al. (2022).

#### Library Preparation and Sequencing

A total of 1,060□ng of extracted DNA was used to prepare a sequencing library with the PacBio SMRTbell prep kit 3.0, following the manufacturer’s instructions (PacBio, Menlo Park, USA). The finished library was sequenced on a PacBio Revio instrument using one SMRT Cell 25M, according to the manufacturer’s recommended protocols. Sequencing was carried out in high-fidelity (HiFi) mode, in which circular consensus sequencing (CCS) is used to generate highly accurate long reads through multiple passes of the polymerase around each SMRTbell template. Raw signal data were processed using PacBio’s integrated CCS and basecalling pipeline to produce reads suitable for downstream genome-level analyses.

### Assembly, Polishing, and Contamination Screening

#### Genome assembly

The PacBio HiFi-reads were assembled using hifiasm v0.19.6 (Cheng et al. 2021, 2022) and flye v2.8.3. Contigs were assigned to different genomes after blast (Camacho et al. 2009) searches, rRNA identification (barrnap, https://github.com/tseemann/barrnap) and organelle assembly. Assembly statistics (assembly size, N50, GC%, contig count) were computed using the tool ‘assembly-stats’.

#### Comparison of the Completeness of the C. muelleri Genome

Genome completeness of the *C. muelleri* genome was evaluated with BUSCO v5.7.1 using the stramenopiles_odb10 lineage dataset (also cross-checked with eukaryota_odb10 for robustness). We reported the percentages of complete (single-copy and duplicated), fragmented, and missing BUSCOs under default scoring.

### Orthologous Group Patterns and Functional Innovation in C. muelleri

#### Proteome Preparation

Gene prediction for *C.*□*muelleri* was performed using the MAKER v3. Annotation pipeline, which integrated evidence-guided training of AUGUSTUS to refine ab initio gene models. To ensure a non-redundant gene set, only one isoform per locus (longest CDS) was retained. Protein models shorter than 50 amino acids or lacking a valid start or stop codon were excluded from downstream analyses.

#### Orthogroup inference across 14 diatom genomes

Reference assemblies and/or published proteomes for 14 centric (5) and pennate (9) diatom species (*Chaetoceros tenuissimus*, *Skeletonema marinoi*, *Thalassiosira pseudonana*, *Thalassiosira oceanica*, *Cylindrotheca closterium*, *Fragilaria crotonensis*, *Fistulifera solaris*, *Fragilariopsis cylindrus*, Mayamaea pseudoterrestris, *Nitzchia inconspicua*, *Phaeodactylum tricornutum*, *Pseudo-nitzschia multistriata*, and *Seminavis robusta* ,Table S3) were compiled using identical parameters to minimize pipeline bias. Where multiple annotations existed, the most recent, well-curated release was used, and the same isoform filtering was applied. OGs were inferred with OrthoFinder v2.5.5 (DIAMOND mode ultra-sensitive), using default MCL inflation (1.5) and automatic rooting. The species tree was reconstructed with STAG and rooted with STRIDE (as implemented in OrthoFinder). OrthoFinder’s duplication mapping was used to place gene duplication events onto the species tree. Species-specific OGs were defined as OGs with members from exactly one species. For *C.*□*muelleri* expansions, an OG was considered expanded if its *C.*□*muelleri* copy number exceeded that of all other species in the 14-genome panel. Also, at the *Chaetoceros* genus level, pair-exclusive OGs were defined as OGs with members present in both *Chaetoceros* species (*C.*□*muelleri*, *C.*□*tenuissimus*) but absent from all other taxa.

### Functional Enrichment of Species-Specific and Expanded Gene Sets

#### Functional Annotation

Functional annotation of the *C.*□*muelleri* proteome were performed exclusively with eggNOG-mapper v2.1.13 (DIAMOND mode, taxonomic scope set to eukaryotes/stramenopiles) using the eggNOG 5.0 orthology database. Additional homology support was obtained by BLASTP v2.7.1 against a UniProtKB diatom protein dataset downloaded on 11□March□2025 (including unreviewed/TrEMBL entries).

Gene Ontology (GO) terms, KEGG Orthology (KO) identifiers, and COG functional categories were all derived directly from eggNOG-mapper annotations. KO identifiers was used to map proteins to KEGG pathways for enrichment analyses.

PFAM domain annotations was obtained from eggNOG-mapper, which integrates precomputed PFAM 32.0 assignments within the eggNOG 5.0 database. Domain frequencies were summarized across the annotated proteome by counting unique occurrences per protein (collapsing multiple hits on the same gene). High-frequency domain families were defined as those with ≥□30 copies, with additional reporting of families with ≥□20 copies to capture moderately abundant domains. These summaries were used to characterize major functional groups (e.g. repeat-containing proteins, kinases, transporters, helicases,) and to highlight retrotransposon-associated PFAM families (e.g., RVT_1, RVT_2, rve, zf-CCHC) indicative of TE-linked genomic expansion.

### Repeat Content

#### De Novo Repeat Library Construction and Masking

A species-specific repeat library was generated with RepeatModeler2 v2.0.3. The library was curated by removing known protein-coding sequences via BLASTP v2.7.1 against the UniProt database, followed by ProtExcluder v1.2. The assembly was masked using RepeatMasker v4.1.2-p1 with the curated library.

### Comparative Genomics of C. muelleri Across 14 Diatom Genomes

#### Genome/Proteome Curation

Reference genomes and proteomes for 14 diatom species (Table□S3) were obtained from NCBI, EMBL-EBI, and DDBJ. For each species, a single representative isoform per gene (longest CDS) were retained, protein sequences shorter than 50□amino acids were removed, and FASTA headers were standardized to ensure unique gene identifiers. See section on OG inference across 14 diatom genomes for additional details.

#### Orthogroup Clustering, and Core/Near-Core Definitions

OGs and counts were inferred with OrthoFinder v2.5.5. “Core OGs” were defined as OGs present in all 14 species; “strict single-copy OGs” were core OGs with exactly one gene per species; and “near-core OGs” were present in 13 of 14 species. Species-specific OGs were present in exactly one species. Lineage sharing (centrics-only, pennates-only, both) for *Chaetoceros* proteomes were quantified by intersecting OG membership across phylogenetic partitions.

### Comparison of C. muelleri and C. tenuissimus

#### Gene Family Copy-Number Asymmetry and Duplication Counts

For each OG, per-species gene copies “expanded in *C.*□*muelleri*” (or *C.*□*tenuissimus*) was defined as copy number strictly greater than that of all other species in the 14-genome panel. Global duplication and species-specific duplication events were identified using OrthoFinder and mapped to the species tree. Duplication counts were normalized by the total protein count of each species’ and by the total number of OGs containing at least one gene from that species to account for variation in the proteome size and OG membership..

#### Enrichment Analysis of C. tenuissimus-Biased OGs

To evaluate functions over-represented among OGs with higher copy number in *C.*□*tenuissimus*, the corresponding *C.*□*muelleri* genes belonging to those OGs (n□=□1,231; 1,135 annotated) were identified and tested for enrichment across GO, KEGG, COG, and PFAM using Fisher’s exact tests (background□=□8,826 annotated *C.*□*muelleri* genes), with Benjamini–Hochberg FDR correction. Terms with very small, expected counts (<□5) were filtered prior to testing or merged into higher-level parent categories when appropriate.

### Lineage-Structured Gene Duplication Across Diatoms

#### Species-Level Duplication Sets and Pan-Diatom Tests

For each species, genes identified under duplication events by OrthoFinder were compiled (terminal-branch-biased events retained after STAG/STRIDE reconciliation). Within-species enrichment of functional categories (GO biological process and molecular function; KEGG pathways; COG categories) were tested using Fisher’s exact tests with Benjamini–Hochberg FDR correction against the annotated background of that species. For the pan-diatom analysed, fold changes (log2 OR) were aggregated across species by category and summarized across ecological and phylogenetic strata (centric vs. pennate; marine vs. freshwater).

#### Ecological and phylogenetic stratification

Species were assigned to centric/pennate and marine/freshwater categories (Table S3). We estimated the distribution of the enrichment statistics per-species(median OR and interquartile range) for key functional categories (e.g., chromatin organization, signal transduction, carbohydrate metabolism, ion transport, membrane organization, stress response). We report the consistent effect-direction patterns (fold-change >□1) to capture convergent trends where no category passed FDR within species.

## Results

### Genome characterization

Long PacBio reads of the *C. muelleri* BS20 strain yielding 44x coverage were used for the genome assembly. After removing four circular contigs corresponding to bacterial contamination, and the filtering the draft assembly for low coverage contigs, e.g. a *Nanochloropsis* contamination and its organelles, the assembly consisted of 54 contigs with a total assembly length of 43,360,835 bp, a contig N50 of 1.40 Mb, 38.8 % of the total repeat content and 17.86 % of classified transposable elements (Table 1).

**Table 1.**
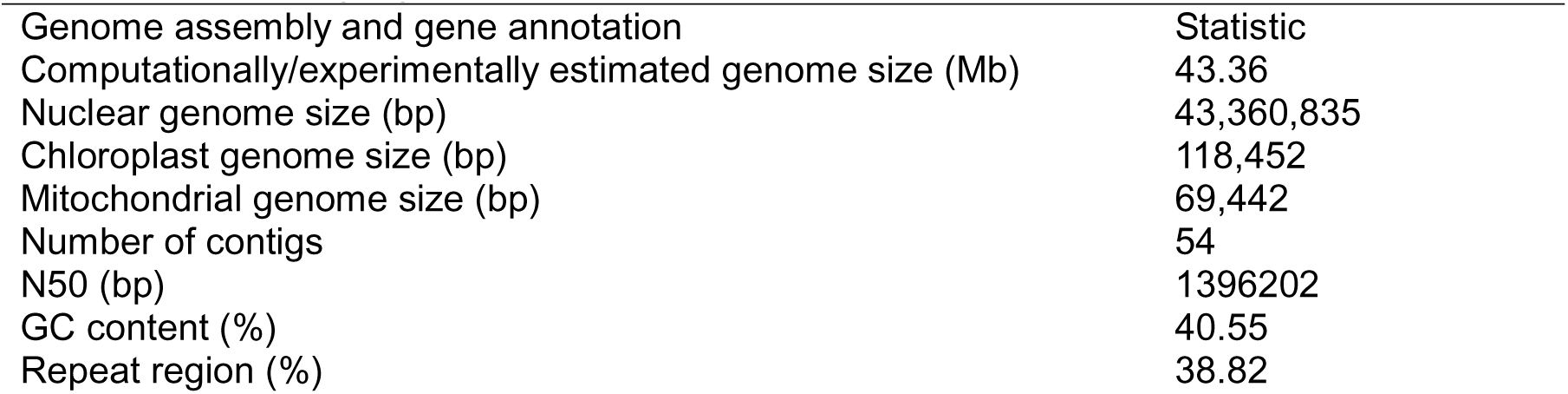
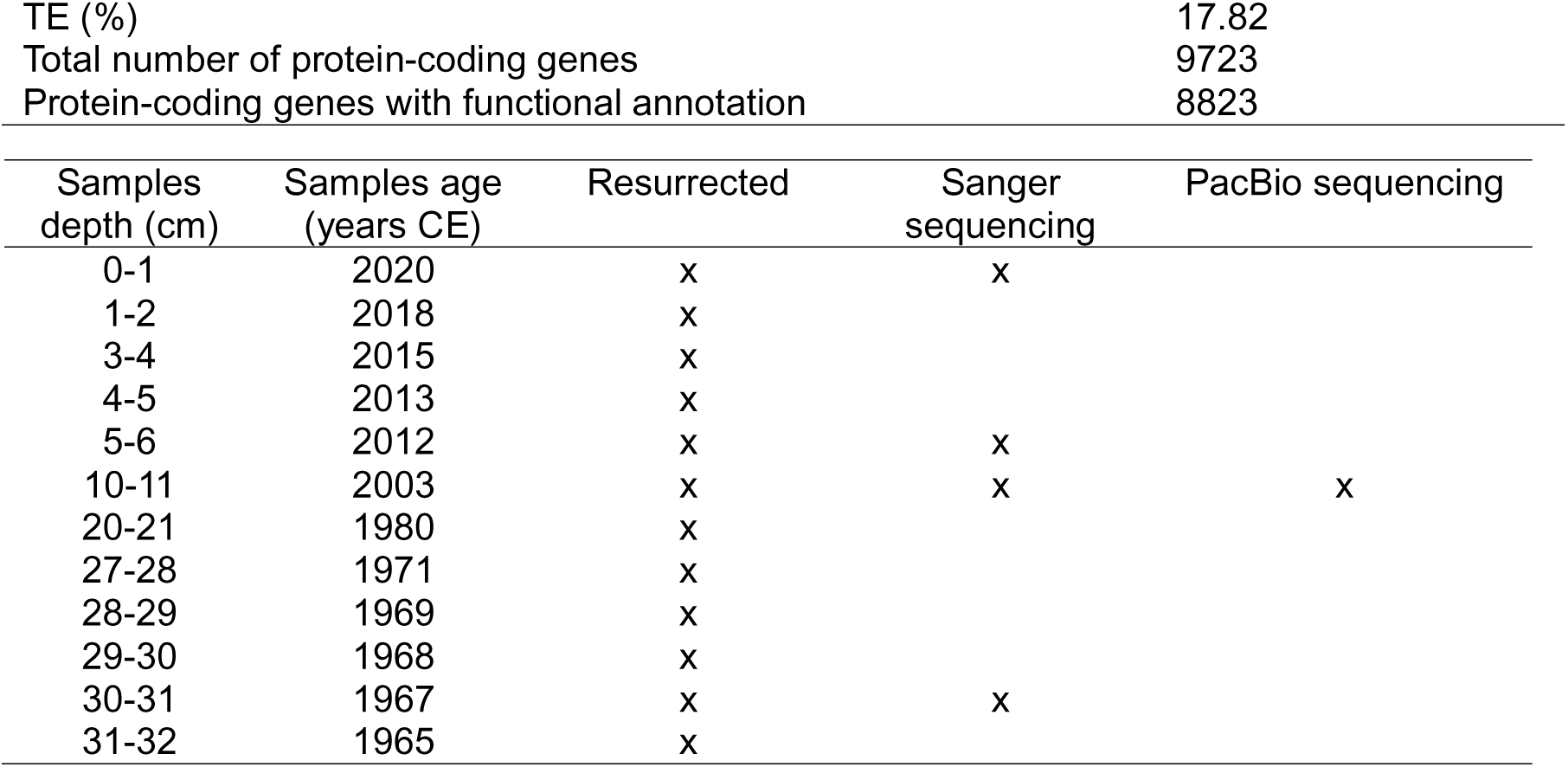
Genome properties of C. muelleri.

| Genome assembly and gene annotation | Statistic |
| --- | --- |
| Computationally/experimentally estimated genome size (Mb) | 43.36 |
| Nuclear genome size (bp) | 43,360,835 |
| Chloroplast genome size (bp) | 118,452 |
| Mitochondrial genome size (bp) | 69,442 |
| Number of contigs | 54 |
| N50 (bp) | 1396202 |
| GC content (%) | 40.55 |
| Repeat region (%) | 38.82 |
| TE (%) | 17.82 |
| Total number of protein-coding genes | 9723 |
| Protein-coding genes with functional annotation | 8823 |

| Samples depth (cm) | Samples age (years CE) | Resurrected | Sanger sequencing | PacBio sequencing |
| --- | --- | --- | --- | --- |
| 0-1 | 2020 | x | x |  |
| 1-2 | 2018 | x |  |  |
| 3-4 | 2015 | x |  |  |
| 4-5 | 2013 | x |  |  |
| 5-6 | 2012 | x | x |  |
| 10-11 | 2003 | x | x | x |
| 20-21 | 1980 | x |  |  |
| 27-28 | 1971 | x |  |  |
| 28-29 | 1969 | x |  |  |
| 29-30 | 1968 | x |  |  |
| 30-31 | 1967 | x | x |  |
| 31-32 | 1965 | x |  |  |

### Comparison of the completeness of the genome

BUSCO analysis indicates that the genome assembly is highly complete, with 93% of the expected orthologous genes identified as complete. Among these, 88% are single copy, suggesting the assembly is largely non-redundant and accurately represents the gene composition. The presence of 5% duplicated BUSCOs may indicate minor assembly redundancy or biological gene duplications, which could affect gene family evolution or copy number studies. Only 1% of BUSCOs are fragmented and 6% are missing, reflecting minimal gaps, robust core gene recovery, and strong assembly continuity) but modestly lower completeness than highly curated model diatoms (*P. tricornutum*, *T. pseudonana*, 97%), and *S. marinoi*, *C. tenuissimus* and *F. cylindrus* also showed strong completeness (≥ 95%, respectively), while *T. oceanica* and *Pseudo-nitzschia multistriata* displayed reduced completeness (90% and 86%, respectively) and higher missing BUSCOs (4–13%), consistent with older assemblies or less accurate annotations. An outlier, *F. solaris*, scored 97% complete yet showed an exceptionally high duplication rate (83% duplicated BUSCOs), aligning with its known allopolyploid genome structure. Overall, these results underscore the high quality of most diatom references while highlighting species-specific differences driven by assembly contiguity, annotation pipelines, and biological complexity (Table S4).

### Orthologous group patterns and functional Innovation in C. muelleri

OG clustering across 14 diatom genomes identified 7,612 OGs containing *C. muelleri* genes. Of these, 156 OGs were specific to *C. muelleri* (i.e., OGs present exclusively in *C. muelleri* and absent from all other species in the 14-genome comparison), comprising 239 species-specific genes. A further 120 OGs were significantly expanded in *C. muelleri* (i.e., OGs found in multiple species but containing a higher copy number in *C. muelleri* than in all other taxa), incorporating 477 genes with *C. muelleri*-specific copy number expansion. These categories, together with the defined background gene universe of 8,823 genes, were used for the enrichment analyses.

### Functional enrichment of species-specific and expanded gene sets

Among the species-specific genes, 119 of 239 genes had GO annotations, while 396 of the 477 expanded gene copies had GO annotations. Enrichment analysis revealed consistent overrepresentation of cell wall biosynthesis, polysaccharide metabolic pathways, and vesicle-mediated trafficking, indicating strong lineage-specific specialization in these processes.

Within the Biological Process category, the most significant enriched terms (q = 7.48×10⁻□) included *plant-type cell wall cellulose metabolic process* (GO:0052541), *cellulose biosynthetic process* (GO:0030244), *plant-type cell wall cellulose biosynthetic process* (GO:0052324), *plant-type cell wall organization* (GO:0009664), and β*-glucan biosynthetic process* (GO:0051274). These processes reflect targeted expansion and diversification of polysaccharide biosynthesis pathways. Broader carbohydrate-related terms—such as *cell wall polysaccharide biosynthetic process* (GO:0070592), *regulation of polysaccharide biosynthetic process* (GO:0032885), and *regulation of carbohydrate metabolic process* (GO:0006109)—were significantly enriched, emphasizing regulatory elaboration alongside enzymatic expansion.

In the Cellular Component category (q = 1.54×10⁻□), enriched terms pointed to intracellular trafficking, secretion, and membrane-derived compartments, including the *trans-Golgi network* (GO:0005802), *cell wall* (GO:0005618), *endosome* (GO:0005768), *Golgi apparatus subcompartment* (GO:0098791), and multiple vesicle-related terms. Thus, these results indicate that *C. muelleri* has evolved an expanded cellular machinery for cell wall construction and vesicle-mediated export of polysaccharide and silica-associated components.

One Molecular Function term was enriched: *identical protein binding* (GO:0042802, q = 0.00339), suggesting selective expansion of proteins involved in homomeric interactions, possibly contributing to the diversification of regulatory or structural complexes (Table S5).

### KEGG pathway enrichment

KEGG analysis (FDR ≤ 0.05) further identified three pathways uniquely enriched in *C. muelleri* relative to 13 reference diatom genomes: glycerophospholipid metabolism (map/ko00564) indicating membrane remodelling innovations; phosphonate and phosphinate metabolism (map/ko00440) suggesting expanded capacity for organophosphorus utilization, and basal transcription factors (map/ko03022) implying diversification in core transcriptional machinery. These pathways complement GO results by highlighting lipid, phosphorus, and regulatory specialization specific to this lineage.

### COG category enrichment and evidence of genome remodeling

Across both species-specific OGs and expanded OGs, COG L (Replication, recombination and repair) was strongly enriched: unique OGs: OR□=□9.85, FDR□=□3.7×10⁻¹□, expanded OGs: OR□=□7.63, FDR□=□1.5×10⁻□□, and pairwise comparison with *C. tenuissimus* revealed: OR□=□2.27, FDR□=□4.2×10⁻□. This pervasive enrichment of DNA metabolism and repair functions is consistent with a genome influenced by extensive TE activity. Further, in eukaryotes, TE-linked proteins are typically classified under COG L rather than the bacterial-oriented “Mobilome” category (COG X), aligning with the observed patterns (Fig. 2).

**Fig.2a.**
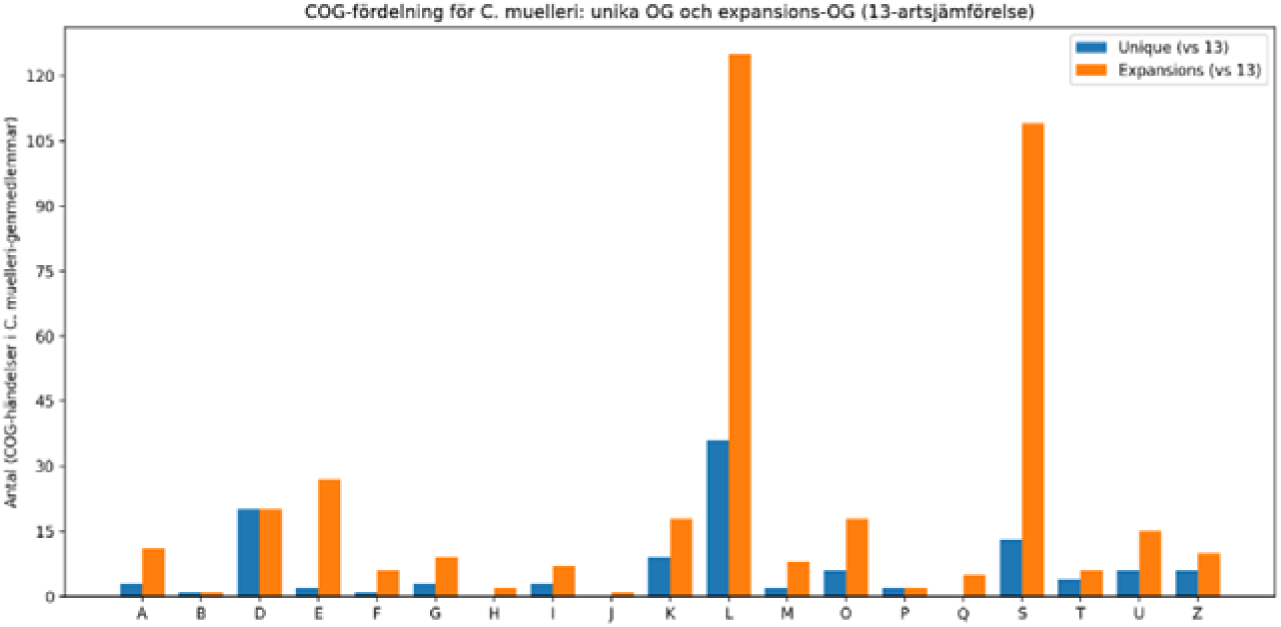
Comparison of unique versus expansions of COG categories in *C. muelleri*

**Fig.2b.**
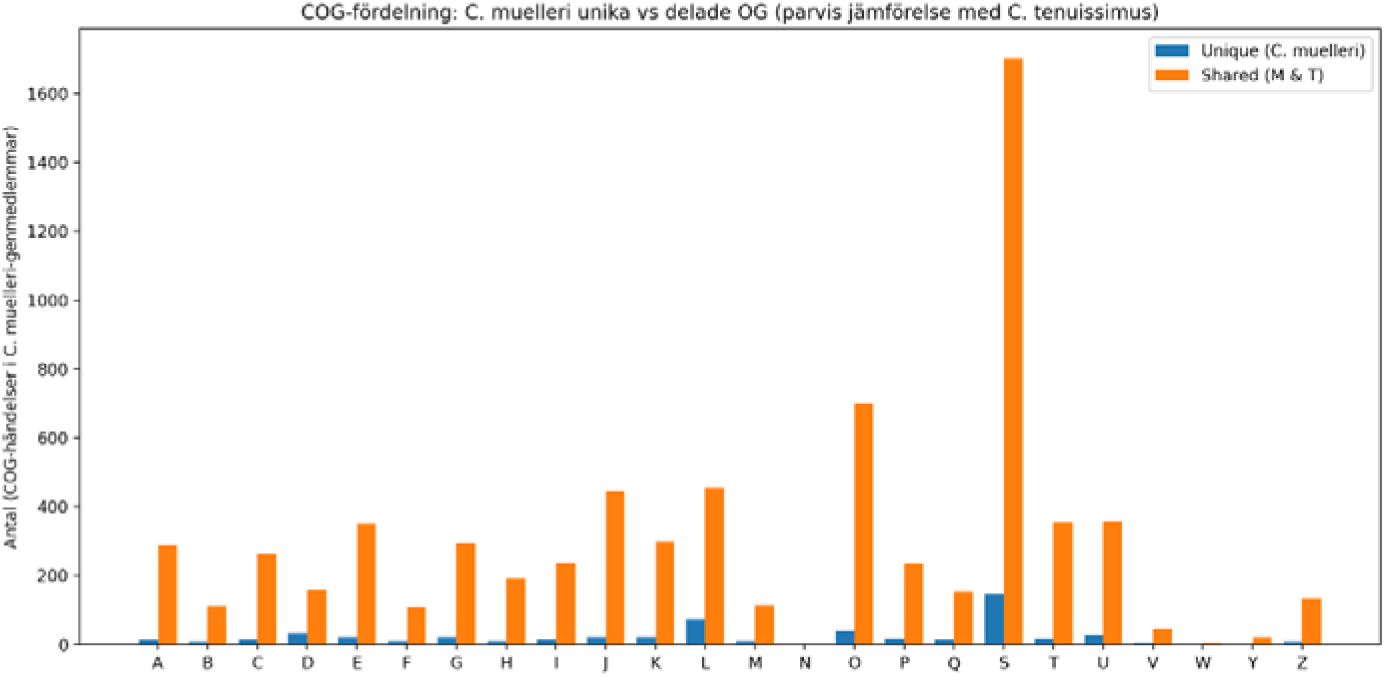
Pairwise comparison of unique versus shared COG categories in *C. muelleri* and *C. tenuissimus*.

### PFAM domain architecture of the C. muelleri genome

PFAM domains were quantified across all predicted gene models to characterize the protein domain architecture of the *C. muelleri* genome. Whole-genome profiling revealed a mixture of deeply conserved eukaryotic protein families and lineage-specific expansions. Across the complete proteome, 31 PFAM domains occurred at high frequency (≥30 copies), representing core functional categories such as protein kinases (Pkinase; 124 copies), RNA/DNA helicases (Helicase_C; 105 copies), WD40 repeat proteins (93 copies), DnaJ-class molecular chaperones (49 copies), and multiple ankyrin repeat subfamilies (ANK, Ank_2, Ank_5). Transport-associated domains were also well represented, including the mitochondrial carrier family (Mito_carr), ATP-binding cassette transporters (ABC_tran), and the major facilitator superfamily (MFS_1). Several domains related to retrotransposon activity exhibited high abundance, most notably the reverse transcriptase families RVT_2 (87 copies) and RVT_1 (41 copies), integrase-associated rve domains (73 copies), and retroelement-linked zinc-finger motifs (zf-CCHC). Lowering the abundance threshold to ≥20 copies expanded the catalog to 58 PFAM domains, revealing moderately frequent categories that included chromatin remodelling modules (SNF2_N), diverse signalling components, short-chain dehydrogenases (adh_short), metal-binding or catalytic motifs (Metallophos, Thioredoxin), and a variety of repeat-containing proteins. Hence, these patterns highlight a proteome shaped by both conserved eukaryotic functions and marked lineage-specific expansion of retrotransposon-derived domains, reflecting the combined influence of core cellular processes and genome-reshaping TE activity on the evolution of the *C. muelleri* gene repertoire. These PFAM patterns highlight the combination of conserved eukaryotic machinery (kinases, helicases, WD40, transporters, metabolic enzymes) and lineage-specific proliferation of retrotransposon-derived domains shape the *C. muelleri* gene repertoire (Table S6).

### Repeat content

Repeat content analysis revealed that approximately 38.82% (16,710,566 bp) of the *C. muelleri* genome (43.05 Mb) consists of repetitive sequences, based on RepeatMasker annotations. TEs account for about 17.86 % (7,687,730 bp) of the genome, dominated by LTR retrotransposons (14.9%), followed by DNA transposons (2.6%), LINEs (0.34%), and Helitrons (0.04%), while SINEs were not detected. Simple repeats and low-complexity regions contribute an additional ∼2.2%, and a large fraction (∼19.3%) remains classified as “Unknown,” indicating potential novel or poorly characterized repeats. (Table 2, Figs. 3 -4).

**Fig. 3.**
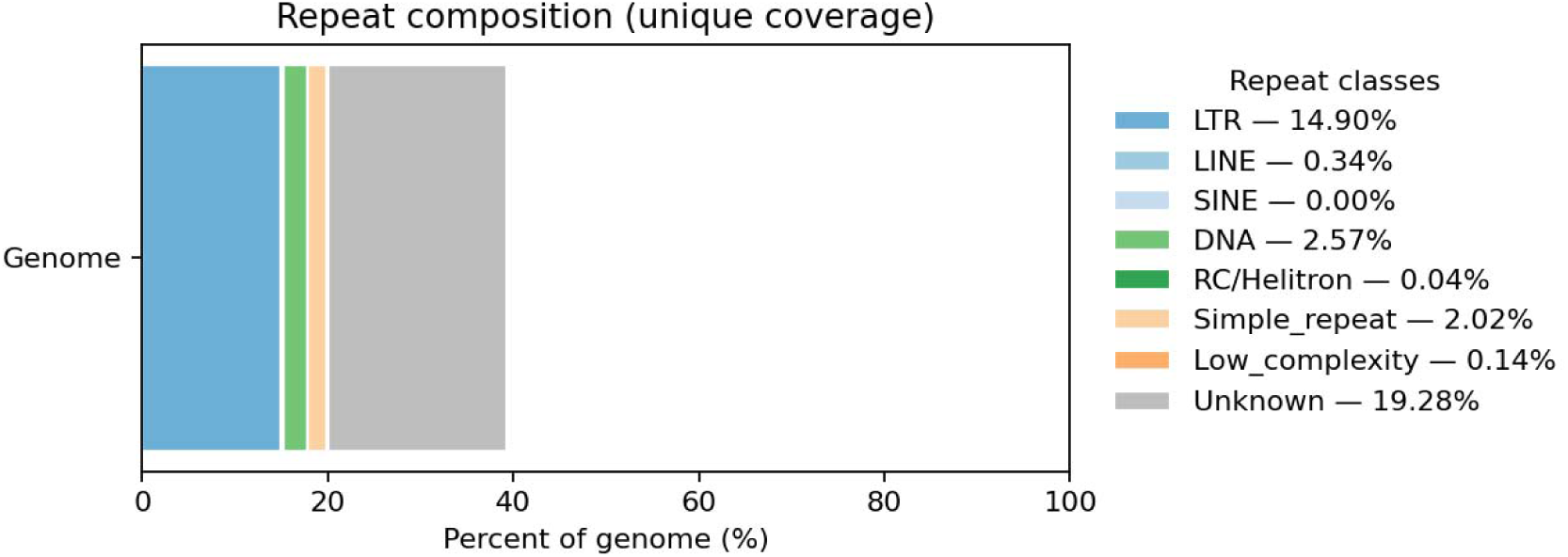
Overall repeats of all the classes of the *C. muelleri* genome

**Fig. 4.**
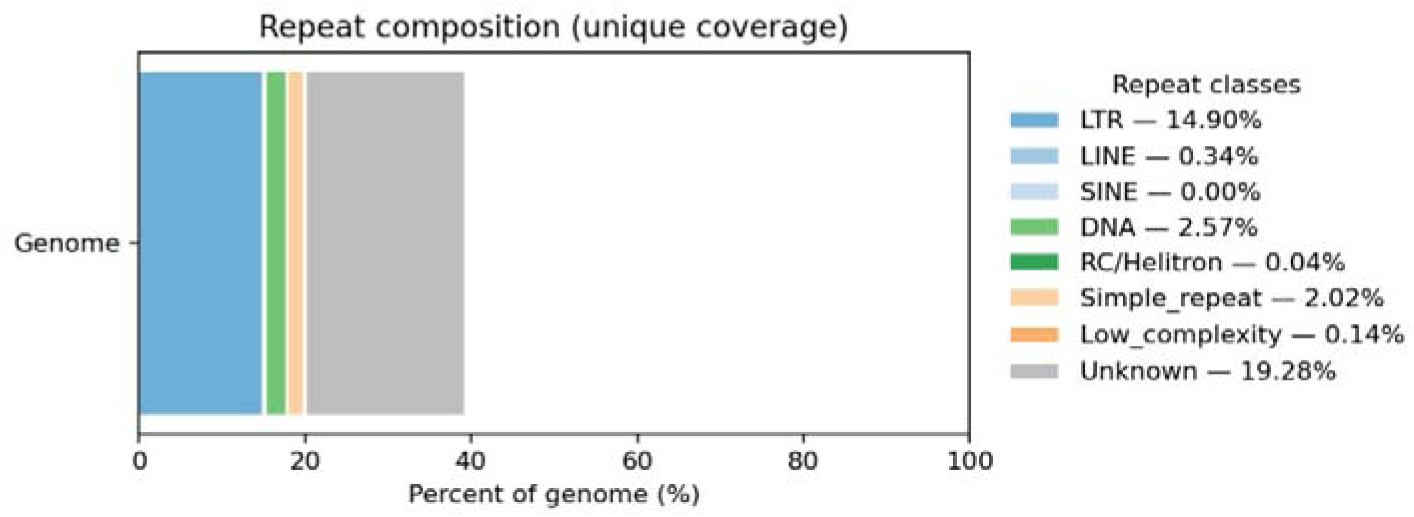
TE landscape in *C. muelleri*

**Table 2.** Superfamilies of repeats.

| genus | n_features | total_bp | mean_bp | median_bp | genome_pct |
| --- | --- | --- | --- | --- | --- |
| Unknown | 4760 | 8,299,763 | 1,743.65 | 359 | 19.2789 |
| Simple_repeat | 166,842 | 8,122,406 | 48.68 | 37 | 18.8669 |
| LTR/Copia | 1,529 | 3,828,265 | 2,503.77 | 956 | 8.8924 |
| LTR/Gypsy | 366 | 1,352,479 | 3,695.30 | 2,146 | 3.1416 |
| LTR | 148 | 1,131,875 | 7,647.80 | 6,334 | 2.6291 |
| Low_complexity | 11,374 | 561,667 | 49.38 | 42 | 1.3047 |
| DNA/PIF-Harbinger | 381 | 374,086 | 981.85 | 855 | 0.8689 |
| DNA/MULE-MuDR | 98 | 230,722 | 2,354.31 | 1,688 | 0.5359 |
| DNA/PiggyBac | 67 | 133,428 | 1,991.46 | 1,690 | 0.3099 |

### Comparative genomics of C. muelleri across 14 diatom genomes

We analyzed 14 well-annotated diatom genomes (Table S3) using OrthoFinder, clustering 267, 195 genes into 27,815 OGs and reconstructing a rooted species tree with duplication event mapping. The global OG landscape comprised 2,583 core OGs (9.3%) shared by all species and only 2 strict single-copy OGs, with a further 1,909 near-core OGs present in 13 of 14 genomes, underscoring extensive paralogy across diatoms. Across the 14-species matrix, species-specific OGs (present in exactly one genome) were abundant (∼38.7% of all OGs; 10,773 OGs total), corresponding to ∼20.1% of all genes (53,645), consistent with widespread lineage-specific innovation.

Species-specific OG counts varied among taxa. In pennate diatoms, values ranged from 85 in *Pseudo-nitzschia multistriata* to 2,209 in *Seminavis robusta*, while centric diatoms spanned 116 in *Thalassiosira pseudonana* to 1,875 in *T. oceanica*, indicating variability both within and across lineages. No significant difference (p> 0,05) in total species-specific OG counts was detected between centric and pennate diatoms. Within *Chaetoceros*, *C. muelleri* exhibited fewer species-specific OGs than *C. tenuissimus* (e.g., 156 vs 702), with the identification of pair-exclusive OGs (present in both *Chaetoceros* species but absent elsewhere (e.g., 51 OGs) (Table S7).

We examined lineage sharing (centrics vs pennates) for the *Chaetoceros* proteomes. Of the 7,612 OGs containing *C. muelleri* genes, 100 were shared exclusively with pennates, 511 exclusively with centrics, 6,845 with both lineages, and 156 were species-specific to *C. muelleri*. For *C. tenuissimus* (10,148 OGs), 591 were pennate-only, 726 centric-only, 8,064 shared with both, and 702 were species-specific. These categories delineate lineage-shared versus species-unique components and suggest further investigation of transport, remodelling, and stress-associated genes.

### Gene-family expansions associated with C. tenuissimus

*C. muelleri* proteins were assigned to 17,655 OGs spanning conserved and lineage-specific functions. Of these, 71.0% (12,527) mapped to diatom-related lineages (Bacillariophyta, *T. pseudonana*, *Phaeodactylum tricornutum*), and 29.0% (5,128) to higher-level eukaryotic groups with conserved domains and occasional bacterial/viral motifs. Within the diatom subset, the largest overlaps were Bacillariophyta–*Thalassiosira* (5,111 groups) and Bacillariophyta–*Phaeodactylum* (3,658), with 828 groups shared by all three. Since eggNOG groups are not identical to OrthoFinder OGs, these results are presented for complementary context.

### Comparison of C. muelleri and C. tenuissimus

Comparison of *the two Chaetoceros* genomes revealed that the gene family copy-number asymmetries were substantial: 548 OGs were expanded in *C. muelleri* whereas 834 OGs were expanded in *C. tenuissimus*, indicating divergent trajectories of gene family amplification. A total of ∼106, 315 duplications were mapped on the species tree, with a strong excess on terminal branches. *C. muelleri* harbored ∼902 high-confidence duplications spanning ∼602 OGs (support□=□1.0). After normalizing duplication counts by proteome size and OG occupancy, *C. muelleri* ranked among the duplication-rich lineages (Table S8), with expansions concentrated in families associated with stress response, signalling, and metabolic flexibility.

To investigate gene-family dynamics between *C.*□*tenuissimus* and *C.*□*muelleri*, we identified OGs in which *C.*□*tenuissimus* harbored a greater number of gene copies than *C.*□*muelleri*. We identified 1,231 *C.*□*muelleri* genes belonging to OGs showing higher copy-number in *C.*□*tenuissimus*. Of these, 1,135 genes carried functional annotations (background n□=□8,826) and were used for enrichment tests across GO, COG, KEGG, and PFAM using Fisher’s exact tests with FDR correction.

The strongest GO enrichments clustered around nucleosome/chromatin organization, led by GO:0006334 (overlap□=□9/11, OR□=□30.7, FDR□=□1.45×10⁻□), GO:0000786 (7/7, OR□=□∞, FDR□=□1.45×10⁻□), GO:0031492 (7/7, OR□=□∞, FDR□=□1.45×10⁻□), and several related terms (e.g., GO:0031490, GO:0031497, GO:0044815). COG categories showed a significant enrichment for P (Inorganic ion transport and metabolism) (61/250, OR□=□2.25, FDR□=□7.0×10⁻□), with other categories not passing FDR. KEGG pathway analysis was dominated by chromatin-heavy maps (e.g., ko05322, overlap□=□16/26, OR□=□11.0, FDR□=□9.35×10⁻□), alongside stress/cell-death signaling (e.g., ko04217). PFAM domain enrichment highlighted HSF_DNA-bind (25/33, OR□=□21.6, FDR□=□1.26×10⁻¹□), Histone (13/17, OR□=□22.3, FDR□=□5.35×10⁻□), Sulfatase/Sulfotransfer_2 (16/17 and 10/11, FDR ≤□1.16×10⁻□), and TE-associated domains (RVT_1 and rve; FDR ≤□1.14×10⁻□), together with Silic_transp (5/5, OR□=□∞, FDR□=□1.40×10⁻□), consistent with an interplay of chromatin remodeling, ion/solute transport, sulfur metabolism, TE activity, and silica cell-wall–linked transport in the *C.*□*tenuissimus*-biased OGs. We note this is a conservative projection based on *C.*□*muelleri* annotations; a symmetric analysis using *C.*□*tenuissimus* gene-wise annotations will refine organism-specific functions in the *C.*□*tenuissimus*-only families (Fig. 5).

**Fig. 5.**
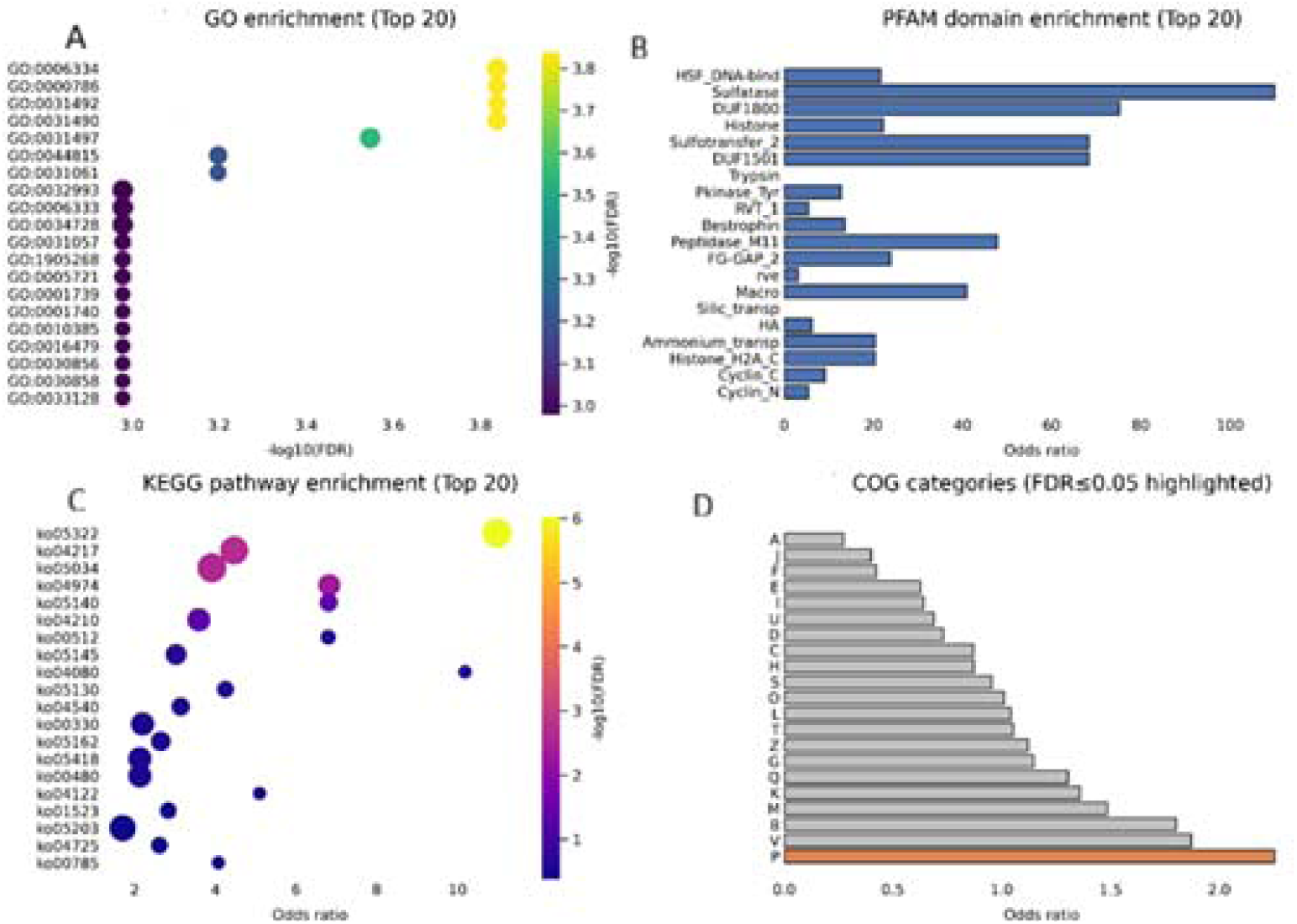
Proxy enrichment for *C. tenuissimus* vs.LJ*C. muelleri* orthogroups A (GO enrichment): Significantly enriched GO terms (Top-20 by FDR) in the *C.*□*muelleri* proxy gene set drawn from orthogroups where *C.*□*tenuissimus* has higher copy-number than *C.*□*muelleri*. Dot size reflects overlap; color encodes −log□□(FDR). B (PFAM): PFAM domain enrichment (Top-20) shown as horizontal bars (odds ratio). Enriched domains include chromatin- and stress-linked modules (e.g., HSF_DNA-bind, Histone), transporters (Bestrophin, Silic_transp), sulfur metabolism (Sulfatase/Sulfotransfer_2), and TE-associated signatures (RVT_1, rve). C (KEGG): KEGG pathway enrichment (Top-20) with bubble size = overlap and color = −log□□(FDR); chromatin-heavy and stress/cell-death signaling maps dominate. D (bCOG): COG category enrichment (bars = odds ratio; orange indicates FDR ≤ 0.05). COG P (Inorganic ion transport & metabolism) is significantly enriched. Design: Target set = 1,135 annotated *C.*□*muelleri* genes from 834 Cten□>□CM orthogroups; background = all annotated *C.*□□*muelleri* genes (n ≈ 8,826). Tests used Fisher’s exact with Benjamini–Hochberg FDR. (Counts from OrthoFinder “Counts” and Orthogroups membership; annotations from eggNOG-mapper.)

### Lineage-Structured Gene Duplication Across Diatoms

Gene duplication analysis across the 14 genomes revealed that the greatest number of duplications in *N. inconspicua* (25, 862) and the least number of duplications in *P. triocrnutum* (2, 546, Table S8). Across the 14 diatom genomes gene duplication reveals a consistent bias towards regulatory and cellular response functions with global (pan-diatom) enrichment tests showing elevated fold changes in RNA-binding, signal transduction, and cell-cycle pathways. However, no categories were significant at FDR threshold (q□≤□0.05), reflecting the statistical dilution that arises when pooling large, functionally heterogeneous OGs. The fold enrichment patterns emphasize that diatom gene family expansion predominantly targets RNA regulatory systems, cell-cycle and signalling pathways, and core informational machinery.

Species-specific duplications across the 14 diatom genomes revealed that the gene duplication and functional expansion patterns showed clear phylogenetic and ecological structuring. Centric species consistently exhibited enrichment in duplicated genes associated with chromatin organization, transcriptional regulation, and carbohydrate metabolism, while pennate species displayed broader expansions in signal transduction, ion transport, membrane organization, and stress-response pathways. Marine taxa showed expansion of stress-signalling genes, osmotic regulation pathways, and metabolic flexibility, whereas freshwater species exhibited stronger duplication in chromatin remodelling and transcriptional control.

Among the centrics, *C. muelleri* showed strong expansion in vesicle-associated functions (GO:0005794), developmental and regulatory pathways (GO:0045595), and KEGG metabolic modules linked to carbohydrate processing (ko00010) and glycosphingolipid biosynthesis (ko00680). *C. tenuissimus* similarly showed duplication of regulatory and developmental pathways (GO:0051240; GO:0045595) and enrichment in amino-sugar metabolism (ko00480). *Skeletonema marinoi* exhibited duplication in chromatin regulation (GO:0031497; GO:0006323) and signal-transduction pathways (ko04350). *T. oceanica* and *T. pseudonana* showed enrichment in chromatin components (GO:0000791; GO:0031490) and duplication of signalling and metabolic pathways (ko05143, ko04972) consistent with open-ocean environmental pressures.

Marine pennate diatoms showed even more pronounced expansion in signalling and adaptive pathways. *F. cylindrus* displayed strong enrichment for transcription factor activity (GO:0001046, GO:0031490) and environmental response (GO:0051703), together with the expansion of KEGG immune-like or defense related pathways (ko05418; ko05145). *Cylindrotheca closterium* exhibited duplication in ion transport and membrane remodelling functions (GO:0006813; GO:0015079) and KEGG lysosomal/apoptotic modules (ko04217). *P. multistriata* showed duplication related to chromatin remodelling (GO:0006323) and carbohydrate metabolism (ko00500). *Nitzschia inconspicua* revealed expanded methyltransferase and nucleotide metabolism functions (GO:0008168; GO:0009165), along with KEGG central metabolic modules (ko01110; ko01100). P*. tricornutum* displayed enriched duplication in transcriptional control (GO:0051094) and stress response (GO:0006979), with KEGG enrichment in regulatory pathways (ko04261; ko05034). *S. robusta* showed expansion in membrane organization and morphogenesis (GO:0005886; GO:0090066), with KEGG enrichment in transporter systems and metabolic processes (ko02010; ko00910).

Freshwater pennate species exhibited distinct regulatory expansion profiles. *Fragilaria crotonensis* showed duplication in transcriptional regulation (GO:0051240; GO:0051094) and enrichment in glucose-regulation pathways (ko04974). *Mayamaea pseudoterrestris* displayed strong chromatin-associated expansion (GO:0000791; GO:0031060) and KEGG enrichment in MAPK and cell-death-related pathways (ko04024; ko04261) (Fig. 6).

**Fig. 6.**
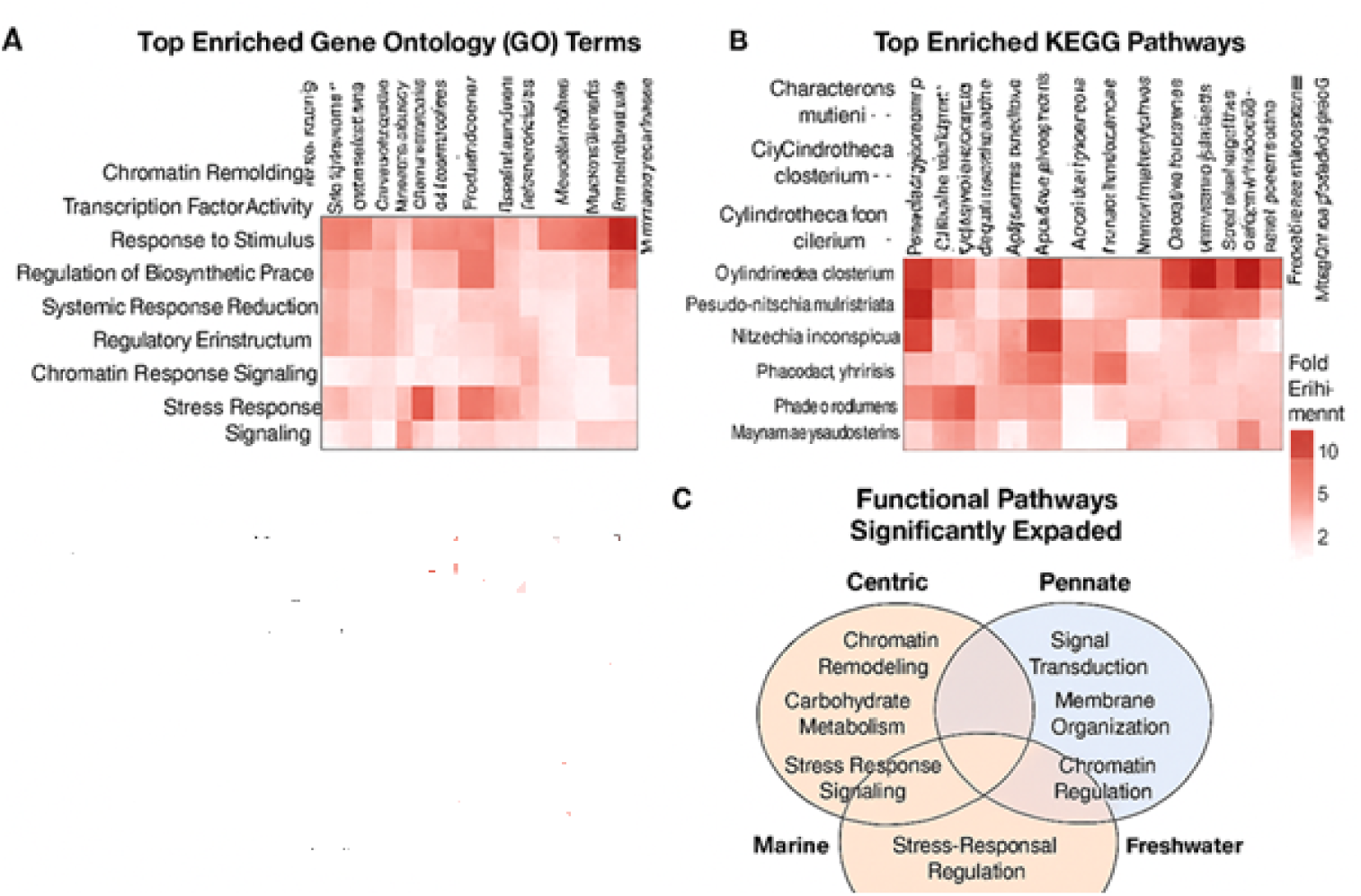
Lineage-Structured Gene Duplication Across Diatoms Heatmap showing presence–absence heatmap of OGs across the 14 genomes. Cells reflect the number of OGs shared by each species pair. Species are ordered by morphology (centric vs pennate), revealing clusters of high OG sharing and broader evolutionary structure. **(A) Top enriched Gene Ontology (GO) terms across species.** Heatmap showing fold enrichment of the most significantly overrepresented GO categories among duplicated or expanded gene families in each genome. Enriched functions include chromatin remodeling, transcription factor activity, response to stimulus, regulatory processes, and stress-response signaling. Values reflect fold enrichment derived from species-specific GO enrichment analyses, highlighting substantial variation in functional expansion patterns across diatom lineages. **(B) Top enriched KEGG pathways across species (first panel).** Heatmap summarizing fold enrichment of significantly overrepresented KEGG pathways associated with duplicated or expanded gene families. Species exhibit distinct expansion signatures across pathways including carbohydrate metabolism (e.g., starch and sucrose metabolism), amino-sugar and nucleotide-sugar metabolism, apoptosis and lysosomal pathways, central carbon metabolism, immune-analog signaling, oxidative phosphorylation, and various stress-response or regulatory modules. These enrichments reflect both lineage-specific evolutionary trajectories and ecological specialization. **(C) Ecological summary of major functional expansions.** Venn-style conceptual diagram integrating phylogenetic (centric vs. pennate) and ecological (marine vs. freshwater) contrasts in functional pathway expansion. Centric species primarily expand chromatin remodeling, carbohydrate metabolism, and stress-response signaling. Pennate species exhibit broader expansion in signal transduction, membrane organization, and chromatin regulation. Marine taxa show strong enrichment in stress-response signaling and transcriptional regulation, whereas freshwater species emphasize chromatin remodeling and transcriptional control. These patterns collectively illustrate how both evolutionary lineage and ecological habitat shape genome-scale functional diversification in diatoms.

Thus, these results demonstrate that gene duplication in diatoms is shaped by both evolutionary lineage and habitat. Centric species are enriched in chromatin and carbon-allocation processes; marine pennates diversify stress-signalling, membrane systems, and metabolic pathways; and freshwater pennates show strong regulatory and chromatin-based expansions consistent with fluctuating osmotic and nutrient environments.

## Discussion

### A compact yet structurally dynamic genome

The compact (43□Mb), highly contiguous (N50□=□1.40□Mb) genome of *C.*□*muelleri* shows high completeness (93% BUSCO) and a predominance of single copy orthologs (88%), indicating accurate gene recovery with minimal redundancy. The *C. muelleri* genome combines high-quality assembly features, the genome is enriched in repeats (38.8%), including a substantial TE fraction (17.9%), revealing extensive historical repeat-driven remodelling. Thus, these features suggest that *C.*□*muelleri* possesses a compact yet dynamically reshaped genome that facilitate lineage-specific innovation. Such genomic architecture are increasingly recognized in diatoms as substrates for lineage-specific innovation.

### Lineage-Specific Gene Family Innovation

OG analyses across 14 diatoms revealed strong species specific and expanded gene families in *C. muelleri*, identifying 156 species-specific OGs (239 genes), 120 significantly expanded OGs (477 genes), and a total of 7,612 OGs contained *C. muelleri* genes. Although fewer strict species-specific OGs were identified in *C. muelleri* compared to its congener *C. tenuissimus* (10,148 OGs), this distribution indicates that *C. muelleri* has a distinct genomic repertoire shaped by both the emergence of novel genes and the expansion of pre-existing families, many arising from duplications.. These patterns highlight the rapid evolutionary turnover characteristic of diatoms and underscore the unique functional adaptations of this species.

### Functional enrichment highlights diversification of cell wall and trafficking systems

Both species-specific and expanded OGs revealed that functional enrichment converged on pathways involved in cell wall biosynthesis (including cellulose, β-glucan, and other polysaccharide processes), intracellular trafficking, and vesicle-mediated transport, together with Golgi and endosomal functions. These enriched categories indicate that *C. muelleri* has undergone substantial diversification of gene families central to frustule formation, silica deposition, polysaccharide secretion, and broader interactions with the environment. Such expansions likely reflect ecological specialization and the need for flexible, tightly regulated cell-wall architecture in dynamic marine habitats.

Furthermore, KEGG enrichments in glycerophospholipid, phosphonate or phosphinate metabolism, and basal transcription factors suggest innovations in regulatory and membrane systems that coordinate these dynamic cell wall processes, and nutrient acquisition (especially phosphorous) strategies. These pathways suggest adaptive strategies that may enhance nutrient flexibility, environmental responsiveness, and regulatory complexity. Furthermore, considerable enrichment of COG L (Replication, recombination, repair) indicates increased demand for DNA repair and genome maintenance which is a signature pattern of genomes with substantial TE activity. The genome appears to have undergone extensive TE-mediated restructuring, and selection for robust recombination and DNA repair machinery indicating that TEs are a central evolutionary force in *C. muelleri*. Finally, PFAM domain analysis reveals that the proteome consists of a high copy number of retrotransposon domains (RVT_1, RVT_2, rve, zf-CCHC), large families of kinases, helicases, WD40 repeats, ABC transporters, and MFS transporters, and the expansions in chromatin remodelling proteins, dehydrogenases, and repeat-containing proteins. Thus, the PFAM domain architecture alongside conserved eukaryotic functions reveals extensive TE-derived persistent retrotransposon expansion suggesting widespread TE-mediated innovation.

### Transposable elements as drivers of genome remodelling

High TE content, enrichment of TE-derived PFAM domains, a large fraction of unclassified repeats, and strong enrichment of COG L collectively underscore the central role of TEs in shaping the *C. muelleri* genome. The dominance of LTR retrotransposons (∼15% of the genome), together with extensive reverse transcriptase, integrase, and zinc-finger domains, indicates substantial historical TE activity, while an additional 19.3% of unclassified repeats suggests the presence of novel or highly diverged TE lineages. Enrichment of DNA replication, recombination, and repair functions further points to compensatory genome maintenance in response to TE-driven structural stress. In *C. muelleri*, the emergence of species-exclusive gene families containing regulatory domains (e.g. SNF2_N, DENN, dDENN), transporters, and metabolic enzymes suggests that TE-related structural variation may have served as a substrate for neofunctionalization. A substantial part of the *C. muelleri* genomic repeat landscape is unexplored, suggesting that the species harbours unique TE lineages. Thus, these repeat patterns likely trigger TE-mediated genomic innovation through structural and regulatory variation possibly leading to rapid lineage-specific adaptation in *C. muelleri*, particularly useful in organisms exposed to fluctuating environmental conditions such as marine diatoms.

### Comparison of TE composition in diatoms

Comparative analysis of published diatom genomes shows that TE content varies widely across the lineage. The centric model species *T. pseudonana* possesses an exceptionally compact 34 Mb genome in which repetitive DNA accounts for only ∼3%, indicating very limited TE proliferation (Armbrust et al. 2004). In contrast, the pennate model *P. tricornutum* (27.4 Mb) contains a higher repeat load of ∼12–14%, with clear signatures of recent TE mobilization contributing to genome diversification (Bowler et al. 2008). High repeat accumulation is found in large and complex diatom genomes. *C. cryptica* contains 59% repetitive DNA in its 171 Mb assembly, making it one of the most TE-rich diatoms sequenced to date (Roberts et al. 2020) while the polar diatom *F. cylindrus* harbors more than 35% repeats embedded within a highly heterozygous genome architecture characteristic of cold-adapted species (Mock et al. 2017, Paajanen et al. 2017). In comparison, the *C. muelleri* genome shows intermediate TE abundance. Our high-contiguity resurrected genome assembly contains 15.99 Mb of annotated TEs with overrepresentation of LTR retrotransposons (specifically Copia) and a large fraction of unclassified repeats. Thus, these comparisons place *C. muelleri* above compact low-TE diatoms such as *T. pseudonana* but below the highly repeat-rich genomes of *C. cryptica* and *F. cylindrus*, underscoring the substantial heterogeneity of TE accumulation and genome evolution across diatoms.

### Diatom-wide comparative genomics and evolutionary context

Comparative analyses across 14 diatom genomes place *C. muelleri* within a broader evolutionary framework characterized by high number of paralogs, pronounced species specificity, extensive *C. muelleri*-specific innovation (∼38.7% of OGs), and a small, conserved core genome comprising 2,583 OGs. *C. muelleri* had fewer species-specific OGs than its congener *C. tenuissimus* (e.g.156 vs 702). Gene duplication patterns are significantly structured by phylogeny and habitat: Centrics tend to expand carbohydrate metabolism and chromatin-associated functions. Marine pennates show broader diversification in signaling, ion transport, and membrane systems. Freshwater pennates exhibit stronger expansion in transcriptional regulation and chromatin remodeling. *C. muelleri* largely adheres to centric-typical patterns but also exhibits unique expansions in vesicle trafficking, metabolic pathways, and cell wall-associated processes, reflecting its own ecological trajectory within this landscape of niche-linked genomic diversification. Diatom genomes are highly dynamic, and *C. muelleri* is relatively conservative compared with *C. tenuissimus*. Yet it still displays numerous lineage-specific innovations, indicating ongoing divergence within *Chaetoceros*.

### Divergent evolutionary pathways within Chaetoceros

Direct comparison with *C. tenuissimus* reveals contrasting strategies of genome evolution, despite shared ancestry. *C. tenuissimus* shows a greater gene family expansion of chromatin, ion transport, stress signaling, sulfur metabolism, and silica-related transport functions, whereas *C. muelleri* shows expansions in gene families involved in polysaccharide biosynthesis, vesicle trafficking, and membrane-associated regulatory modules. Furthermore, TE-associated domains are enriched in both species, suggesting that similar underlying mechanisms—repeat mobilization and repair-mediated duplication likely leading toward distinct functional outcomes.

### Evolutionary and Ecological Implications

Lineage-specific functional expansion coupled with TE-driven genome plasticity has enabled *C. muelleri* to refine systems crucial for frustule formation, extracellular matrix production, and environmental interaction. Frustule architecture and exopolysaccharide secretion directly influence buoyancy regulation, defense, nutrient acquisition, and ecological competitiveness. The large number of expanded gene families associated with wall structure and secretory function suggests that these traits may have been key targets of selection. Additionally, enrichment of membrane remodeling and transcriptional pathways points to deeper regulatory reorganization that may enhance environmental responsiveness.

### Future directions

This study provides a comprehensive genomic and comparative framework, and highlight opportunities for further work. The large “Unknown” repeat fraction implies substantial uncharacterized TE diversity yet to be resolved. Improved repeat annotation using custom diatom libraries, structural TE detection, and phylogenetic classification will likely refine understanding of TE dynamics. Chromosome-scale assemblies and isoform-resolved transcriptomics would clarify the structural context of gene expansions and the regulatory architecture of repeat-derived genes. Functional validation through transcriptomic profiling under environmental stress, targeted gene manipulation, or comparative phenotyping—will be essential for linking expanded gene families to ecological strategies.

### Conclusions

The *C. muelleri* genome illustrates how a compact diatom genome can be shaped by TE-mediated remodeling and lineage-specific functional innovation. Repeat-driven duplication, regulatory rewiring, and expansion of key biosynthetic and secretory pathways have collectively contributed to its unique genomic identity and ecological potential. These findings underscore the central role of genome plasticity in diatom evolution and provide a foundation for deeper mechanistic and ecological studies of cellular innovation in marine phytoplankton.

## Data availability

All sequencing data and genome assemblies generated in this study are publicly available. The *Chaetoceros muelleri* BS genome assembly, raw long-read and short-read sequencing datasets, and RepeatMasker annotations will be deposited in the NCBI Sequence Read Archive (SRA) and NCBI Genome under the BioProject accession. The previously published *C. muelleri* reference genome used for comparative analyses is available through NCBI under accession GCA_ _019693545.1. Scripts used for genome assembly are archived on GitHub at https://github.com/tseemann/barrnap) Any additional data supporting the findings of this study are provided within the article, or are available from the corresponding author upon reasonable request.

## Author contributions

Anushree Sanyal designed the experiment, generated and analyzed the data and wrote the manuscript. Elinor Andrén did the field work, sampling, analyzed the data and wrote the manuscript. Christian Tellgren-Roth analyzed the data and wrote the manuscript.

## Competing interests

The authors declare no conflicts of interest.

## Acknowledgements

We thank the captain and crew of R/V Electra af Askö for their excellent assistance with coring operations despite pandemic-related restrictions and challenging weather conditions. We gratefully acknowledge Prof. P.□G. Appleby and G.-T. Piliposian (University of Liverpool) for the rapid processing of the radiometric dating analyses. We thank Anna Vilaplana Burgos for her assistance with the DNA extraction. Funding was provided through the REVIVE project, supported by the Foundation for Baltic and East European Studies (grant no.□42-19). This work was supported by the National Bioinformatics Infrastructure Sweden (NBIS) at SciLifeLab, Sweden. The computations were performed on resources provided by the Swedish National Infrastructure for Computing (SNIC), partially funded by the Swedish Research Council through access to the Dardel supercomputer at the PDC Center for High Performance Computing, KTH Royal Institute of Technology under Project SNIC 2022-22-9. PacBio sequencing was performed by the National Genomics Infrastructure (NGI) Uppsala, SciLifeLab, Sweden.

---

Table S1. Resurrection and sequencing data of the samples from the EL20-SH01-01 core.

**Table S2.** Radiometric chronology and loss on ignition data of the Western Gotland Basin core EL20-SH01-01. The 1986 and 1963 marker horizons are identified from the ¹³⁷Cs record, and sediment ages and sedimentation rates are derived from ²¹⁰Pb using a piecewise CRS model constrained by the ¹³⁷Cs dates.

| Depth |  | Chronology |  | Sedimentation Rate |  |  |  | Loss on ignition (LOI) |
| --- | --- | --- | --- | --- | --- | --- | --- | --- |
| cm | $\text{g cm}^{-2}$ | Date AD | Age y | ± | $\text{g cm}^{-2} \text{y}^{-1}$ | $\text{cm y}^{-1}$ | ± (%) | % |
| 0.0 | 0.00 | 2020 | 0 | 0 |  |  |  |  |
| 1.5 | 0.06 | 2018 | 2 | 1 | 0.033 | 0.77 | 8.9 | 29.86111 |
| 3.5 | 0.15 | 2015 | 5 | 2 | 0.029 | 0.68 | 9.4 | 30.75219 |
| 7.5 | 0.32 | 2009 | 11 | 2 | 0.028 | 0.63 | 10.2 | 29.56417 |
| 8.5 | 0.37 | 2007 | 13 | 2 | 0.027 | 0.52 | 10.9 | 28.67948 |
| 11.5 | 0.53 | 2002 | 18 | 2 | 0.027 | 0.49 | 12.4 | 28.78117 |
| 12.5 | 0.59 | 1999 | 21 | 2 | 0.021 | 0.34 | 12.7 | 30.7947 |
| 14.5 | 0.72 | 1993 | 27 | 3 | 0.030 | 0.38 | 14.6 | 27.5956 |
| 17.5 | 0.98 | 1986 | 34 | 4 | 0.053 | 0.49 |  | 17.32579 |
| 20.5 | 1.37 | 1980 | 40 | 4 | 0.083 | 0.69 |  | 27.14799 |
| 23.5 | 1.71 | 1977 | 43 | 5 | 0.081 | 0.77 |  | 21.82834 |
| 26.5 | 2.00 | 1972 | 48 | 5 | 0.059 | 0.68 |  | 28.44295 |
| 29.5 | 2.23 | 1968 | 52 | 6 | 0.058 | 0.63 |  | 24.63357 |
| 32.5 | 2.55 | 1963 | 57 | 6 | 0.071 | 0.54 |  | 20.66496 |
| 35.5 | 3.01 | 1957 | 63 |  | 0.14 | 0.66 |  | 13.58856 |
| 38.5 | 3.83 | 1954 | 66 |  | 0.38 | 1.14 |  | 6.948597 |

**Table S3.** Diatom genomes included in the comparative genomics analyses.

| Species | Genome assembly accession number | Annotation source |
| --- | --- | --- |
| Centric |  |  |
| *Chaetoceros tenuissimus* | GCA_021927905.1 | DDBJ |
| *Thalassiosira pseudonana* | GCA_000149405.2 | Genbank |
| *Thalassiosira oceanica* | GCA_000296195.2 | Genbank |
| *Skeletonema marinoi* | GCA_030871285.1 | Genbank |
| Pennate |  |  |
| *Cylindrotheca closterium* | GCA_933822405.4 | EMBL |
| *Fragilaria crotonensis* | GCA_022925895.1 | Genbank |
| *Mayamaea pseudoterrestris* | GCA_027923505.1 | DDBJ |
| *Nitzschia inconspicua* | GCA_019154785.2 | Genbank |
| *Phaeodactylum tricornutum* | GCA_000150955.2 | Genbank |
| *Pseudo-nitzschia multistriata* | GCA_900660405.1 | EMBL |
| *Seminavis robusta* | GCA_903772945.1 | EMBL |
| *Fistulifera solaris* | GCA_002217885.1 | Genbank |
| *Fragilariopsis cylindrus* | GCA_001750085.1 | Genbank |

**Table S4.** Comparison of the BUSCO completeness across species.

| Species | Complete (C) | Fragmented (F) | Missing (M) | Notes |
| --- | --- | --- | --- | --- |
| *Phaeodactylum tricornutum* | 97.0% | 3.0% | 0.0% | UniProt Proteome UP0000007591 |
| *Thalassiosira pseudonana* | 97.0% | 3.0% | 0.0% | UniProt Proteome UP000001449 |
| *Fragilariopsis cylindrus* | 95.0% | 4.0% | 1.0% | UniProt Proteome UP000095751 |
| *Skeletonema marinoi* | 97.0% | 0.0% | 3.0% | Swedish Reference Genome Portal (GCA_030871285.1) |
| *Chaetoceros muelleri* | 93.0% | 1.0% | 6.0% | BUSCO v5.7.1 (proteins) |
| *Thalassiosira oceanica* | 90.0% | 6.0% | 4.0% | UniProt Proteome UP000266841 |
| *Pseudo-nitzschia multistriata* | 86.0% | 1.0% | 13.0% | UniProt Proteome UP000291116 |
| *Chaetoceros tenuissimus* | 95.0% | 0.0% | 5.0% | UniProt Proteome UP001054902 |

**Table S5.**
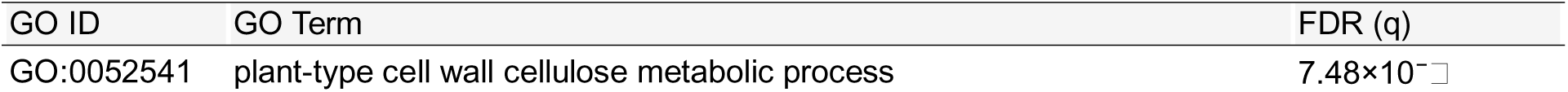
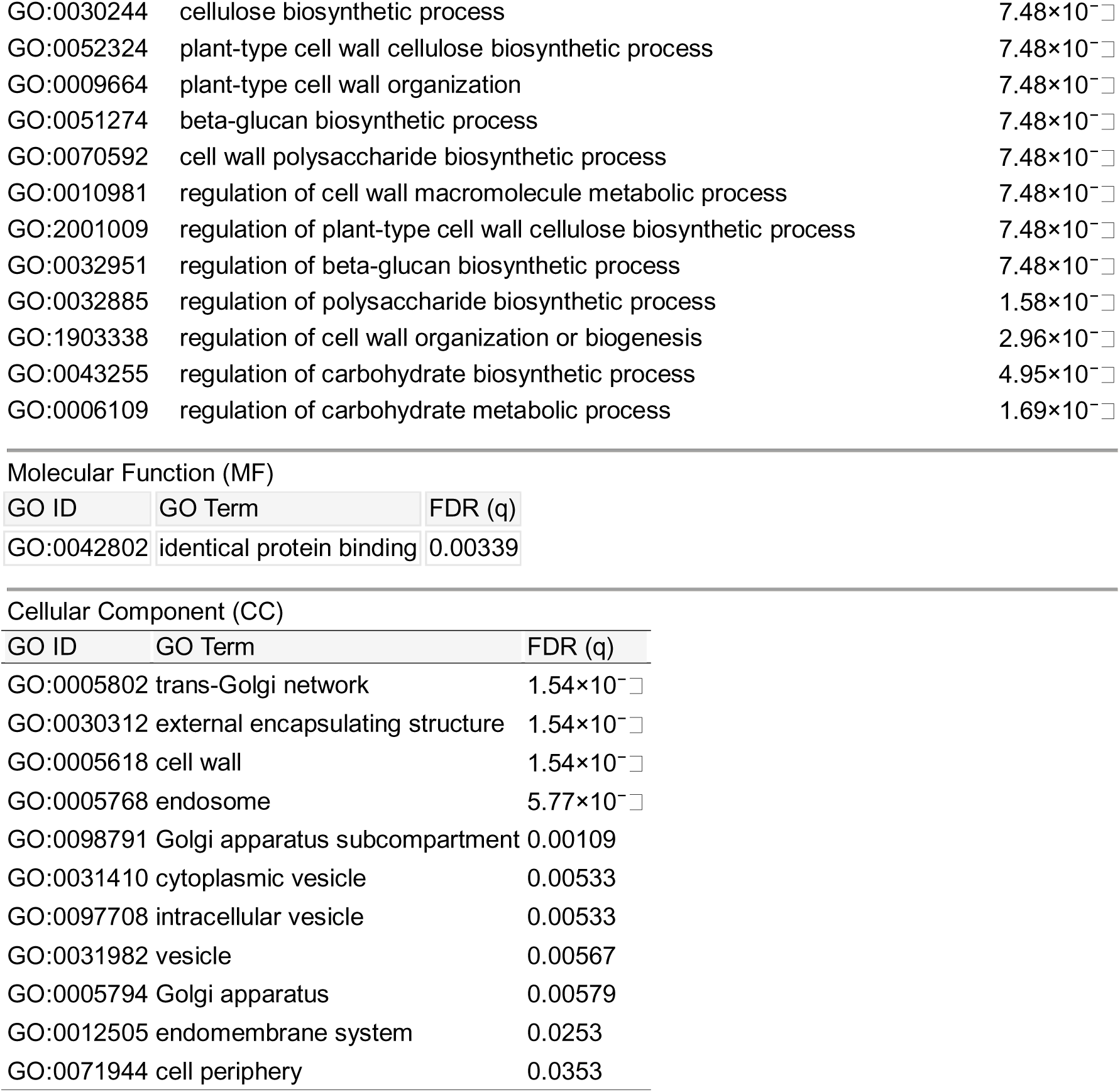
Enrichment of GO terms in the species-specific *C. muelleri* gene set Biological Process (BP)

**Table S6.**
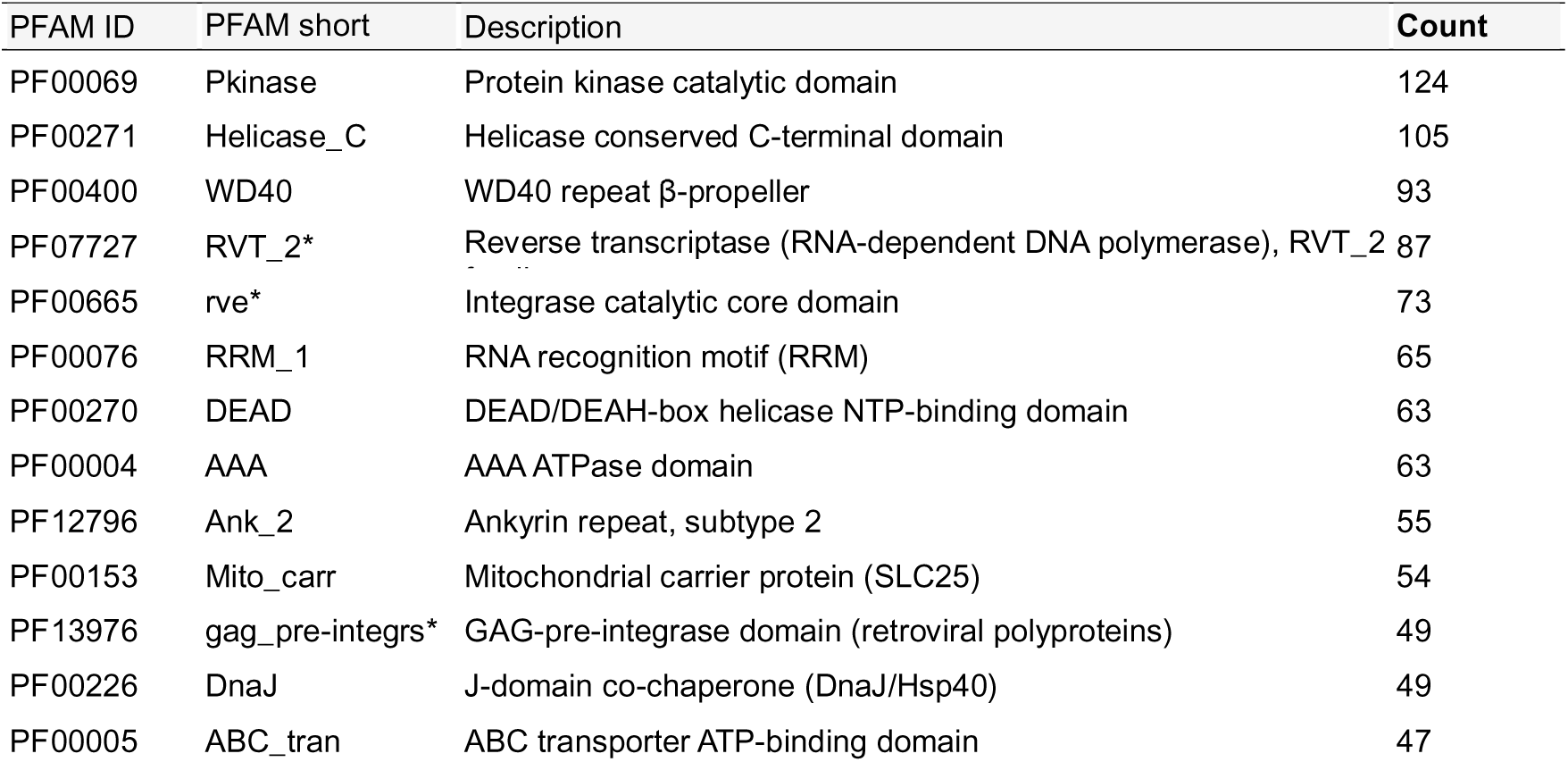
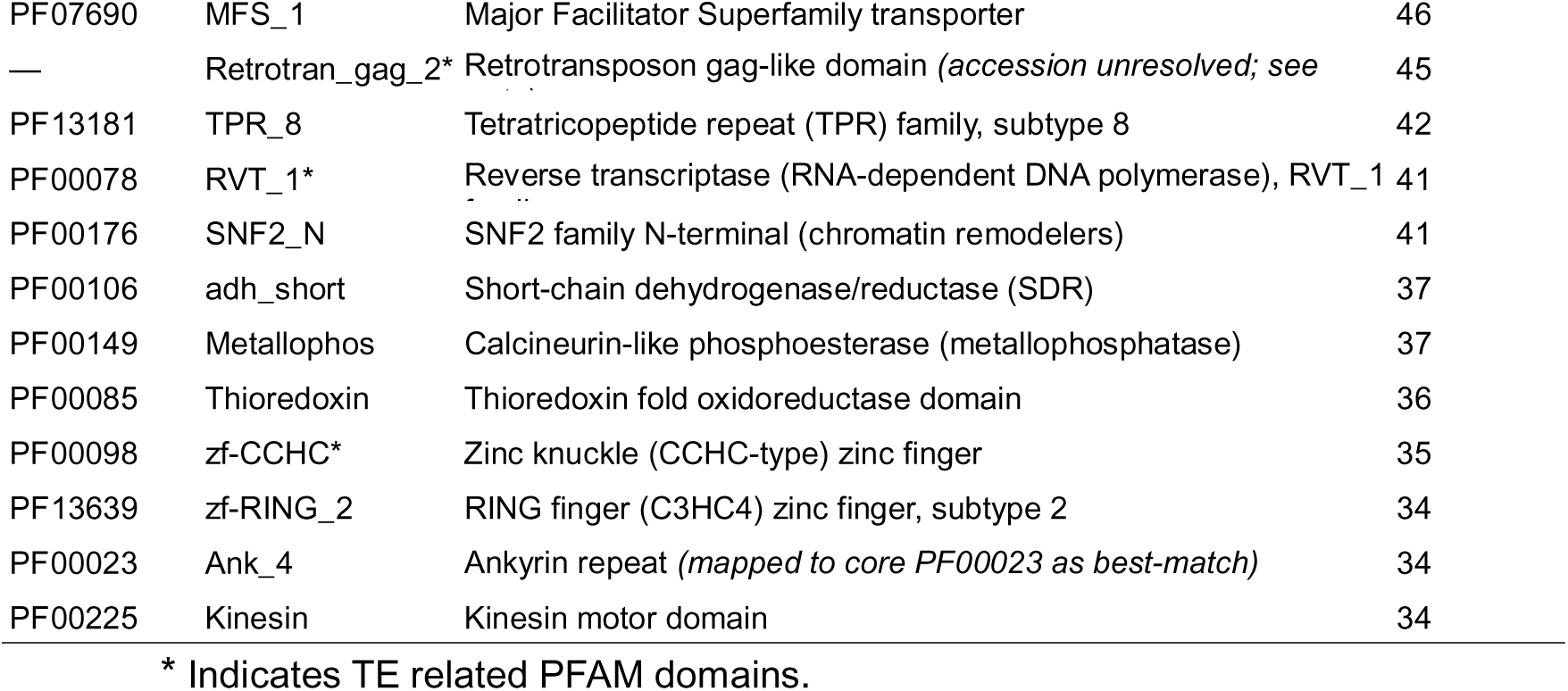
All PFAM domains ≥ 30 hits.

**Table S7.**
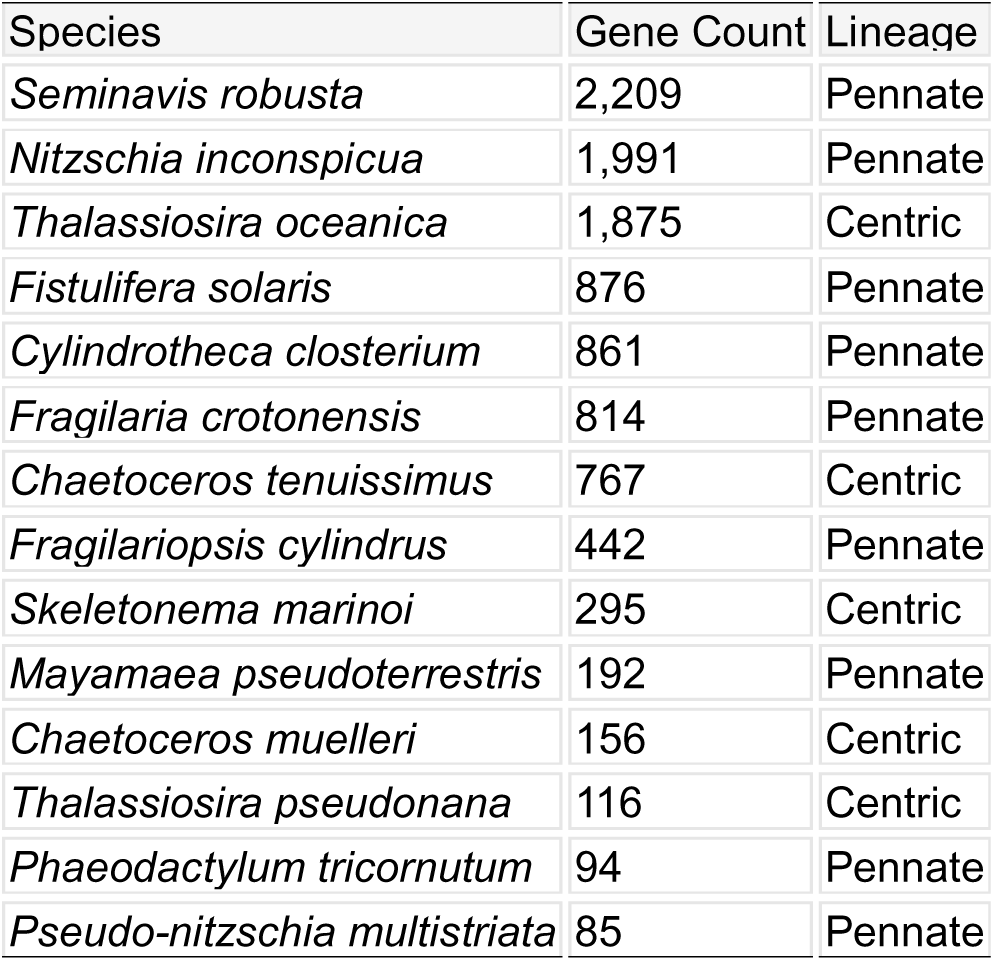
Orthogroups per species with corresponding diatom lineage (centric vs. pennate)

**Table S8.**
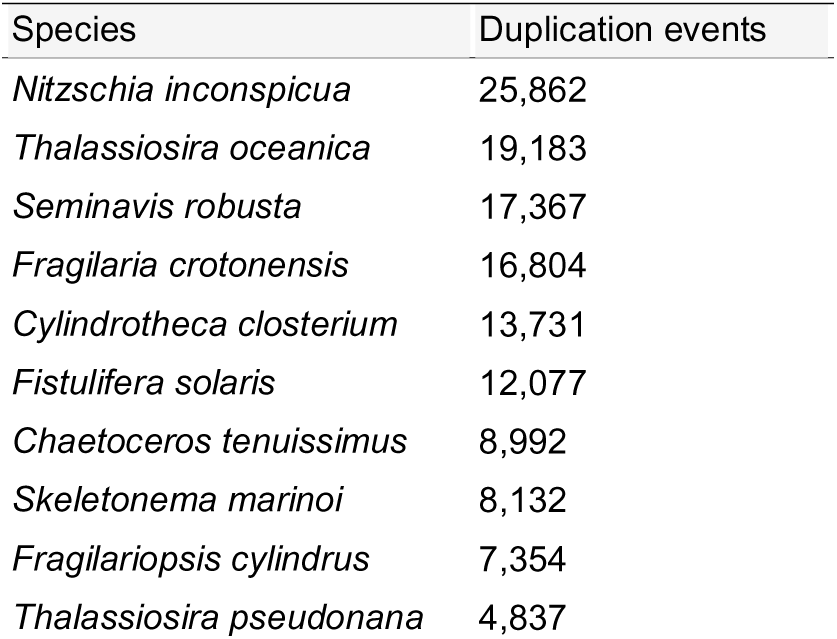
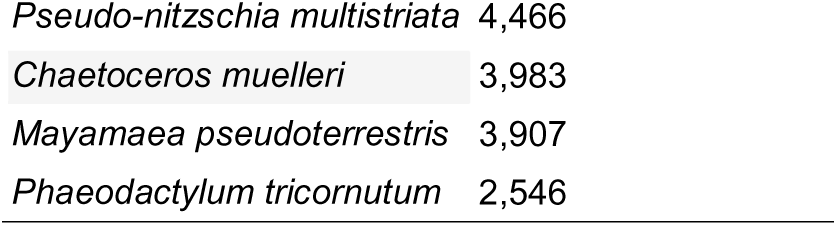
Duplication events in the diatom genomes.

## Notes

### Competing Interest Statement

The authors have declared no competing interest.

